# NMDA receptor hypofunction on GABAergic interneurons results in input-specific Excitatory/Inhibitory imbalance in pyramidal neurons of medial prefrontal cortex

**DOI:** 10.64898/2026.08.12.744479

**Authors:** Carlos A Pretell Annan, Juan E Belforte, Diego E Pafundo

## Abstract

Excitatory/inhibitory (E/I) balance in cortical circuits is typically treated as a global neuronal property, yet pyramidal neurons integrate synaptic inputs from anatomically distinct afferent pathways whose regulation may not be uniform. Using a mouse model with early postnatal NMDA receptor (NMDAR) ablation from corticolimbic GABAergic interneurons with associated E/I dysfunction, we tested whether interneuron NMDAR hypofunction disrupts the mPFC E/I balance globally or in a pathway-specific manner. Here, we use pathway-specific anatomical labeling, optogenetic circuit mapping, paired pyramidal neuron–fast spiking interneuron recordings, and analysis of synaptic integration and found that both structural and functional E/I imbalance emerged selectively at ventral hippocampal (vHPC) inputs onto mPFC pyramidal neurons, while callosal (contralateral mPFC) inputs remained unaffected. Structurally, this imbalance was restricted to vHPC-derived synapses on apical, but not basal, dendrites. Functionally, mutant mice showed an excitation-shifted E/I ratio specific to vHPC-driven responses, arising from a marked failure to recruit feedforward inhibition via fast-spiking interneurons, whose preferential excitatory drive from vHPC inputs was selectively lost. Short-term synaptic plasticity of vHPC and callosal inputs onto both pyramidal neurons and fast spiking interneurons was unchanged, indicating that presynaptic release dynamics could not account for the deficit. Consistent with impaired feedforward inhibition, pharmacological GABA-A receptor blockade failed to prolong vHPC-evoked EPSPs in mutant pyramidal neurons, in contrast to its clear effect on callosal-evoked responses, demonstrating that inhibitory control over the temporal integration of hippocampal, but not callosal, inputs was lost. Altogether, these findings establish pathway-specific E/I imbalance as a consequence of interneuron NMDAR hypofunction, and thus indicate that circuit dysfunction can selectively bias the processing of specific afferent pathways rather than produce a uniform disruption of cortical excitability, with direct relevance for understanding hippocampal–prefrontal dysconnectivity in neuropsychiatric disorders.

## Introduction

The ability of cortical circuits to process and transmit information relies on a precise balance between excitatory and inhibitory synaptic activity. This dynamic equilibrium, commonly referred to as the excitatory/inhibitory (E/I) balance, regulates neuronal excitability, spike timing, synaptic integration, and information flow across cortical networks. Accordingly, disruption of cortical E/I balance has emerged as a common pathophysiological feature of a wide range of neurological and psychiatric disorders, including epilepsy, autism spectrum disorders, schizophrenia, and major depression, although the underlying cellular and circuit mechanisms are likely to differ across these conditions^1–4^. Consistent with this concept, functional E/I balance has been directly quantified and shown to be altered in several animal models of neuropsychiatric disorders^5–11^. Furthermore, experimental manipulation of cortical E/I balance has even been shown to rescue behavioral deficits in animal models, suggesting a causal relation between disease-related behavioral phenotypes and E/I imbalance^12,13^. In parallel, structural analyses in both humans and animal models have revealed corresponding changes in the relative density of excitatory and inhibitory synapses, indicating that functional E/I imbalance is often accompanied by remodeling of synaptic connectivity^14,15^. Together, these findings have established E/I imbalance as a central framework for understanding circuit dysfunction in brain disease^16,17^.

Despite its widespread use, the concept of E/I imbalance has generally been considered a global property of cortical circuits or individual neurons^18,19^. However, cortical pyramidal neurons (PNs) integrate thousands of glutamatergic synapses arising from anatomically and functionally distinct brain regions, while receiving inhibitory inputs from multiple classes of local GABAergic interneurons that differentially target specific subcellular compartments. Moreover, accumulating evidence indicates that synaptic plasticity and circuit remodeling are frequently pathway-specific rather than uniformly distributed across all afferent connections^20–23^. These observations raise the possibility that E/I balance itself may be regulated in an input-specific manner, allowing pathological conditions to selectively disrupt the E/I balance of individual afferent pathways while preserving others.

This possibility is particularly relevant to schizophrenia, a disorder in which converging genetic, pharmacological, and physiological evidence supports a central role for NMDA receptor (NMDAR) hypofunction and impaired GABAergic interneuron function^24–27^. Functional imaging studies demonstrate altered connectivity between the ventral hippocampus (vHPC) and the medial prefrontal cortex (mPFC), a circuit critically involved in working memory, executive function, and cognitive flexibility^28,29^. In parallel, animal models based on developmental NMDAR hypofunction reproduce many behavioral and circuit abnormalities associated with schizophrenia and have repeatedly demonstrated excitation-shifted E/I balance within the prefrontal cortex^30^. Nevertheless, whether these alterations reflect a generalized disturbance of cortical circuitry or preferential dysfunction of specific afferent pathways remains unknown.

We previously generated a mouse model lacking NMDARs selectively in corticolimbic GABAergic interneurons during early postnatal development^31,32^. These mice reproduce multiple behavioral, neurochemical, electrophysiological, and network abnormalities relevant to schizophrenia, including an excitation-shifted structural and functional E/I balance in medial prefrontal pyramidal neurons. However, these previous analyses assessed the integrated activity of all synaptic inputs converging onto individual neurons and therefore could not determine whether the observed imbalance represented a global property of cortical circuits or selectively affected specific afferent pathways.

Here, we tested the hypothesis that interneuron NMDAR hypofunction produces an input-specific rather than generalized disruption of cortical E/I balance. Combining pathway-specific anatomical labeling, optogenetic circuit mapping, paired electrophysiological recordings, and functional analysis of synaptic integration, we demonstrate that structural and functional E/I imbalance selectively affects ventral hippocampal inputs onto medial prefrontal pyramidal neurons, whereas callosal corticocortical inputs remain balanced. Mechanistically, this selective imbalance results from impaired recruitment of feedforward inhibition onto fast-spiking interneurons (FSIs), leading to prolonged temporal integration of hippocampal inputs, but not callosal ones. These findings identify pathway-specific E/I imbalance as a previously unrecognized principle of cortical circuit dysfunction and provide a conceptual framework for understanding how interneuron dysfunction selectively biases the processing of distinct afferent pathways relevant for neuropsychiatric diseases.

## Materials and Methods

A brief description of materials and methods is provided below, full details are available in the Supplementary Materials and Methods.

### Animals

Male and female Grin1 conditional knockout mice (Ppp1r2-Cre - floxed-Grin1; KO) and Cre-negative littermates (control) were used, following protocols approved by the University of Buenos Aires School of Medicine IACUC and government regulations (SENASA, Argentina). In KO mice Cre-mediated deletion of obligatory Grin1 subunit occurs in ∼50% of cortical and hippocampal GABAergic interneurons, including over 75 percent of parvalbumin positive neurons, from the second postnatal week^31,32^.

### Surgery and viral labeling

Mice (8-10 weeks) received stereotaxic microinjections of AAV-CaMKIIa-hChR2(H134R)-EYFP into either the vHPC or the mPFC to enable pathway specific synaptic detection and optogenetic stimulation. In a subset of animals, an AAV PHP.eB -pAAV-CAG-tdTomato adeno-associated virus was additionally injected in the retroorbital sinus to sparsely label mPFC pyramidal neurons for structural analysis. Viral spread was confirmed histologically for each animal.

### Immunohistochemistry

Sparsely tdTomato labeled pyramidal neurons were immunostained for mCherry, VGAT and GFP to visualize pyramidal neurons, and their dendritic segments and dendritic spines together with inhibitory VGAT+ appositions and vHPC or callosal EYFP+ axonal appositions to dendritic spines respectively. Images were acquired by structured-illumination microscopy, and dendritic segments, spines and pucta were reconstructed and quantified in Neurolucida.

### Slice physiology

Whole cell recordings were obtained, 13-14 weeks after viral injection, from mPFC layer 2/3 pyramidal neurons and FSIs in acute coronal slices ipsilateral to vHPC injections and contralateral to mPFC injections. ChR2 expressing vHPC or callosal axons were stimulated with 447nm light to evoke EPSCs, IPSCs and EPSPs. In a subset of experiments, EPSPs were recorded before and after bath application of the GABA-A receptor antagonist picrotoxin (50 μM).

### Quantification and Statistical Analysis

Structural E/I was calculated as the density of excitatory inputs (spine or pathway specific appositions) divided by the density of VGAT appositions to the dendritic shaft. Functional E/I was calculated evoked EPSC area divided by evoked IPSC area ( -60mV and 0mV respectively). Short term depression and paired pulse ratio were derived from trains of optogenetic stimulation^33^. Data are reported as mean ± SEM. Normality was assessed with Shapiro-Wilk tests; group comparisons used paired/unpaired t-tests or ANOVA (with log-transformation when normality was not met), as indicated in the Results and Figure Legends. One neuron was excluded from structural E/I analysis (Figure 1I, Suppl Figure 3D) as a Grubbs-test outlier (α=0.05). Analyses were performed in Prism (GraphPad) and Statistica (StatSoft).

**Figure 1.**
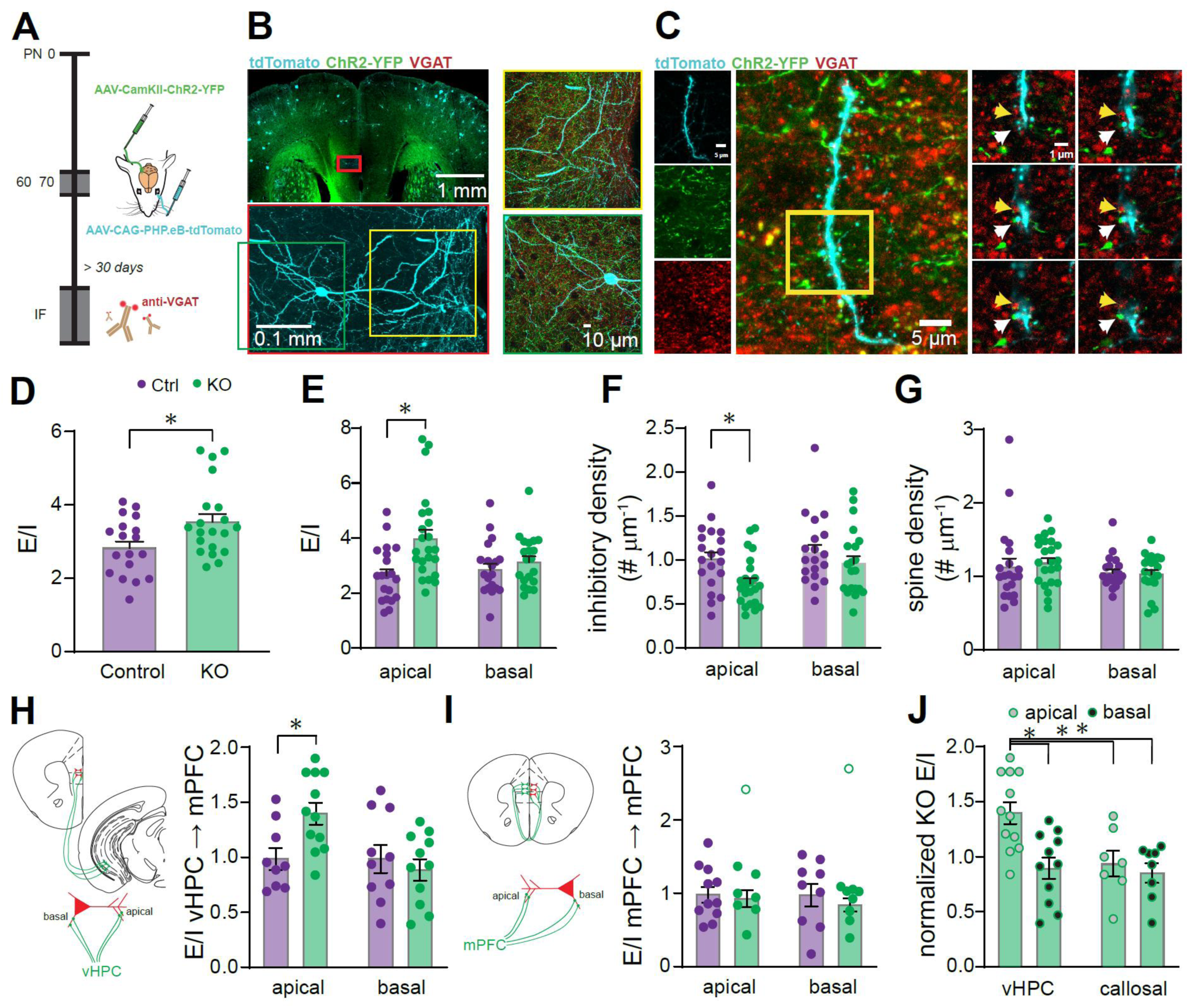
Interneuron NMDAR hypofunction leads to an input-specific structural E/I imbalance in the dendritic tree of mPFC pyramidal neurons. **A**, Schematic depiction of the experimental design. At 60–70 days postnatally, the AAV-CaMKIIa-hChR2(H134R)-EYFP vector was microinjected into either the vHPC or mPFC, and the PHP.eB-AAV-CAG-tdTomato vector was administered intravenously via retro-orbital injection. At 30–50 days post-surgery, animals were perfused, and coronal brain sections were processed for immunofluorescent staining. Inhibitory VGAT-positive appositions onto apical and basal dendrites of sparsely labeled mPFC pyramidal neurons were quantified. In separate experiments, excitatory ipsilateral vHPC or contralateral callosal appositions onto dendritic spines were quantified in the corresponding dendritic segments. **B, top left**, representative image of a coronal section of mPFC of a vHPC microinjected mouse. tdTomato labeled neurons are sparsely distributed throughout the brain including mPFC and green-labeled fibers from the ipsilateral vHPC are present in the left mPFC. The red rectangle indicates the location of the analyzed pyramidal neuron. **Bottom left**, Z-projection of a fluorescence image stack of a representative tdTomato-filled pyramidal neuron. **Right**, zoomed-in images of apical and basal dendrites from the neuron shown on the left. Hippocampal fibers are shown in green, and VGAT-positive puncta are shown in red. **C**, detailed representative image of an analyzed dendritic segment. **Left**, Z-projections of tdTomato-labeled dendritic segments, with EYFP-labeled vHPC axons shown in green and VGAT-positive puncta shown in red. **Middle**, composite Z-projection. **Right**, Individual optical planes from the region demarcated by the yellow rectangle in the middle image, showing a representative VGAT-positive punctum apposed to the dendrite (yellow arrow) and a YFP-labeled bouton apposed to a dendritic spine (white arrow). **D**, global dendritic E/I balance, calculated as the ratio of spine density to VGAT-positive apposition density in dendritic segments, averaged per neuron in control and KO mice (unpaired t-test on log-transformed data, t_(36)_=2.482 p = 0.0179, n = 18–20 neurons). **E**, global dendritic E/I balance calculated across different dendritic segments reveals a compartment-specific E/I imbalance in KO mice, preferentially affecting apical dendrites (mixed-effects ANOVA on log-transformed data: dendrite compartment × genotype interaction, F_(1,38)_= 4.239, p = 0.046 n = 20-23; Sidak’s post hoc test for control vs. KO: apical p = 0.001, basal p = 0.713). **F**, VGAT inhibitory apposition density is decreased in the apical segment of KO mice (mixed-effects ANOVA on log-transformed data: compartment effect, F_(1,37)_= 4.306 p = 0.045 n = 20-23; genotype effect, F_(1,41)_=6.934 p = 0.012; compartment × genotype interaction, F_(1,37)_=1.080 p = 0.305; Sidak’s post hoc test for control vs. KO: apical p = 0.018, basal p = 0.451). **G**, spine density does not change in the KOs in either dendritic segment (mixed-effects ANOVA on log-transformed data: compartment effect, F_(1,38)_= 1.854 p = 0.181 n= 20-23; genotype effect, F_(1,41)_=0.120 p = 0.731; compartment × genotype interaction, F_(1,38)_= 1.324 p = 0.257). **H**, Input-specific excitatory/inhibitory balance calculated as excitatory vHPC appositions onto dendritic spines, normalized to the average control value for each dendritic segment. Hippocampal inputs show an excitation-biased imbalance in the apical sections of KO mice (mixed-effects ANOVA: dendrite compartment × genotype interaction, F_(1,16)_= 6.249 p = 0.024 n = 9-13; Sidak’s post hoc test for control vs. KO: apical p = 0.022, basal p=0.749). I, Input-specific excitatory/inhibitory balance calculated for excitatory callosal appositions onto dendritic spines, normalized to the average control value for each dendritic segment. Callosal inputs show no alteration in E/I balance in KO mice (mixed-effects ANOVA: compartment effect, F_(1,13)_=0.175 p = 0.683 n = 7-11; genotype effect, F_(1,18)_= 0.281 p = 0.603; compartment × genotype interaction, F_(1,13)_= 0.004 p = 0.951; one data point identified as an outlier by Grubbs’ test (α = 0.05, open circle) was excluded from the statistical analyses. J, normalized E/I balance in KO across all experiments reveals a specific imbalance in the apical segment of vHPC inputs. (two-way ANOVA: compartment × input region, F_(1,34)_= 4.135 p = 0.049 n = 7-12; Sidak’s post hoc tests: apical vs basal vHPC p = 0.003, apical vHPC vs apical callosal p= 0.023, apical vHPC vs basal callosal p= 0.003, apical callosal vs basal callosal p = 0.996). In all cases, bar plots show mean ± SEM, with individual values indicated by circles. Purple, control; green, KO mice. *p < 0.05, as indicated in each case.

## Results

Previous reports in schizophrenia patients and in mouse models useful for the study of schizophrenia describe E/I imbalances in the prefrontal cortical circuit both functionally and structurally^2,30,35^. Based on these findings, we initially sought to confirm our previous observations of structural E/I imbalance after ablation of NMDAR early during neurodevelopment in corticolimbical interneurons by quantifying the ratio of global excitatory and inhibitory synaptic inputs onto the dendritic arbor of medial prefrontal cortex pyramidal neurons. To this end, we sparsely labeled pyramidal neurons by retro-orbital injection of a diluted PHP.eB-CAG-tdTomato AAV vector, which is capable of crossing the blood–brain barrier, and visualized GABAergic presynaptic boutons by vesicular GABA transporter (VGAT) immunofluorescence (Figure 1A). Thus, we were able to analyze dendritic segments in the apical and basal dendrites in tdTomato labeled single pyramidal neurons (Figure 1B-C and Suppl Figure 1). We found that within the dendrites, the KO mice show structural E/I imbalance towards excitation between the dendritic spines and GABAergic puncta (Figure 1D) indicating that dendritic integration may be altered in the mouse model. Thus, an important question was whether the structural E/I imbalance is a general feature of KO pyramidal neurons or whether it exhibits spatial selectivity within the dendritic arbor. Compartment-specific analysis revealed that the structural E/I imbalance was restricted to the apical dendrites, with KO mice exhibiting a 35% increase in the E/I ratio compared with controls (mixed-effects ANOVA, dendrite compartment × genotype interaction, p = 0.046; Sidak’s post hoc test: apical, p = 0.001), while the basal dendrites remained unaffected (p = 0.713, Figure 1E). In the KO, the E/I imbalance appears to be the result of a reduction in the GABAergic puncta density in the dendrites (Figure 1F, mixed effects ANOVA; genotype effect, p = 0.012, Sidak’s post-test p=0.018 control vs KO in the apical dendrites and p= 0.451 in the basal dendrites) rather than an increase in the total excitatory inputs (Figure 1G). Whereas GABAergic synapses are predominantly of local origin, glutamatergic synapses onto spines arise from both local pyramidal neurons and long-range projections from distant brain regions. Because different excitatory inputs are known to undergo pathway-specific regulation and plasticity, we hypothesized that the structural E/I imbalance might also be input-specific rather than a global property. To test this hypothesis, we examined two major glutamatergic projections to the mPFC: the ventral hippocampus, a long-range input supporting working memory, recognition memory, goal-directed learning, and executive function^28,36,37^, and the mPFC callosal projection (contralateral mPFC), a cortico-cortical pathway connecting homologous cortical areas at the same hierarchical level^38^. Visual inspection of apical and basal dendrites revealed abundant EYFP-positive puncta apposed to dendritic spines of labeled mPFC pyramidal neurons (Figure 1C and Suppl Figure 1). No significant differences were observed between control and KO mice in the proportion of labeled spines receiving either vHPC or callosal inputs (Suppl Figure 2), indicating that early NMDAR ablation does not alter the relative representation of these afferent inputs onto the dendritic spine population. However, analysis of the structural E/I balance for individual afferent pathways revealed a significant imbalance in KO mice of vHPC inputs restricted to the apical dendritic compartment (Figure 1H and Suppl Figure 3; mixed-effects ANOVA, dendrite compartment × genotype interaction p =0.024; Sidak’s post hoc test: apical dendrites, p = 0.022; basal dendrites, p = 0.749). In contrast, callosal inputs showed no changes in structural E/I balance between control and KO mice, either in apical or basal compartments (Figure 1I and Suppl Figure 3; mixed-effects ANOVA, interaction p =0.951). Furthermore, direct statistical comparison of KO inputs normalized to controls across pathways and dendritic compartments confirmed that the structural E/I imbalance was selectively present at vHPC-derived synapses on apical dendrites (Figure 1J). Together, these findings demonstrate that early NMDAR hypofunction does not produce a uniform global disruption of structural E/I balance, but instead leads to an input-specific structural E/I imbalance that selectively affects ventral hippocampal synapses onto the apical dendrites of mPFC pyramidal neurons.

Functional disruption of hippocampal–prefrontal connectivity has been reported in patients with schizophrenia and in relevant animal models, including mice with interneuron NMDAR hypofunction. Although our previous work demonstrated an excitation-shifted E/I balance in mPFC pyramidal neurons in these mutant mice, the measurements reflected the integrated activity of all synaptic inputs converging onto the recorded cells. Having identified an input-specific structural E/I imbalance, we next asked whether this synaptic alteration is accompanied by selective functional deficits in hippocampal–prefrontal connectivity. To quantify the functional E/I balance of individual afferent pathways onto mPFC pyramidal neurons, we recorded optogenetically evoked excitatory (EPSCs) and inhibitory (IPSCs) synaptic currents from the same neuron while voltage-clamping at −60 and 0 mV, respectively. Channelrhodopsin-2 (ChR2) was selectively expressed in glutamatergic neurons of either the ipsilateral vHPC or, in separate experiments, the contralateral mPFC by local injection of AAV2-CaMKII-hChR2(H134R)-EYFP-WPRE-pA (Figure 2A-C). Whole-cell recordings were obtained from layer 2/3 pyramidal neurons and ChR2-expressing axons surrounding the recorded neurons were stimulated through a water immersion 40× objective centered on the recorded neuron, to determine the relative excitatory and inhibitory synaptic drive provided by each projection (Figure 2D). Optogenetic stimulation of either vHPC or callosal axons within the mPFC evoked short-latency, low-jitter EPSCs at −60 mV, consistent with direct monosynaptic excitation from both pathways (Figure 2E and Suppl Table 1). Next, we generated input–output curves by stimulating each pathway at different light intensities to evaluate the strength of the excitatory drive form both pathways (Figure 2F). This analysis revealed a selective decrease in excitatory synaptic drive onto mPFC pyramidal neurons in KO mice in response to vHPC, but not callosal, stimulation (Figure 2G). In the same neurons, voltage-clamping at 0 mV revealed IPSCs with the expected short delay relative to the EPSCs (4.1–5.3 ms), consistent with feedforward disynaptic inhibition recruited by both projections (Figure 2H and Suppl Table 1). Analysis of the functional E/I ratio revealed an input-specific imbalance following interneuron NMDAR ablation. KO mice exhibited a significant excitation-shifted E/I balance in response to vHPC stimulation, whereas callosal inputs remained unaltered (Figure 2I). Unexpectedly, this shift toward excitation occurred despite the reduced excitatory drive evoked by vHPCinputs, suggesting a marked impairment in inhibitory control. Because both vHPCand callosal projections are glutamatergic, the optogenetically evoked IPSCs necessarily arise through local feedforward inhibitory circuits. Given that parvalbumin-positive fast-spiking interneurons receive direct excitatory input from both pathways and are principal mediators of feedforward inhibition in the mPFC, we next examined their functional recruitment. Consistent with the excitation-shifted E/I balance of vHPC inputs, whole-cell recordings from mPFC fast-spiking interneurons revealed a marked reduction in excitatory synaptic drive from vHPC, but not callosal, inputs in KO mice (Figure 2J).

**Figure 2.**
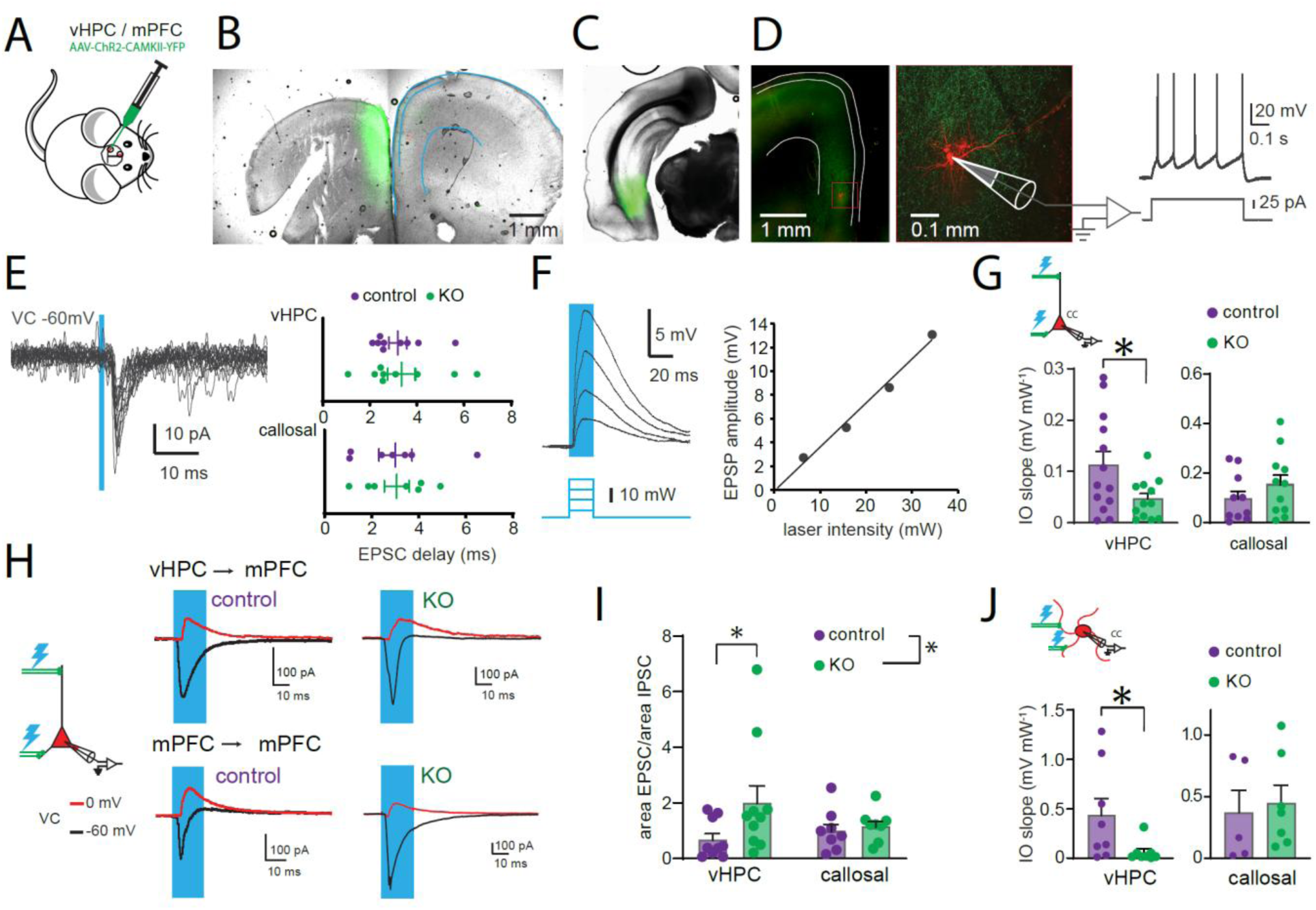
Input-specific functional E/I imbalance of individual afferent pathways onto mPFC pyramidal neurons following interneuron NMDAR ablation. **A,** AAV-CaMKIIa-hChR2(H134R)-EYFP vector was microinjected either in the ventral hippocampus (vHPC) or the mPFC. **B,** representative coronal slice from an mPFC-injected mouse. Green-fluorescence indicates EYFP expression at the injection site including labeled neurons (left hemisphere). In the contralateral hemisphere (right), red cell corresponds to a recorded, neurobiotin-filled neuron and green fluorescence indicates EYFP expression in light-stimulated callosal axons. **C,** Representative coronal slices from a vHPC-injected mouse. Injection site in the left vHPC (green). **D, left,** mPFC of the same mouse as in C, showing a recorded pyramidal neuron (red) and ChR2-containing vHPC axons (green) on the ipsilateral hemisphere. **Middle,** detailed image of the boxed region indicated in the image on the left. Z-projection of a fluorescence stack showing the tdTomato-filled pyramidal neuron (red) and ChR2-containing vHPC axons (green). **Right,** representative voltage response of the same neuron to a depolarizing current step. **E, left,** representative individual sweeps of inward currents evoked at −60 mV by optogenetic activation of ChR2-containing axons (1 ms, 447 nm laser pulse; 20 sweeps) in a pyramidal neuron from a vHPC-injected mouse. **Right,** latency of light-evoked EPSCs in mPFC pyramidal neurons from vHPC-and callosal-injected control and KO mice. **F,** Representative input–output responses of a pyramidal neuron recorded in current-clamp mode to increasing 447 nm laser intensities (20 ms pulses). **G,** Input–output slope of pyramidal neuron responses in vHPC-and callosal-injected control and KO mice. (vHPC: unpaired t-test p=0.035, n=12-13; callosal: unpaired t-test p=0.271, n=10-11;). **H,** EPSCs and IPSCs recorded from the same pyramidal neuron by voltage-clamping at −60 mV and 0 mV, respectively. Representative traces show EPSCs (black) and IPSCs (red) evoked by a 20 ms, 447 nm laser pulse. **I,** functional E/I balance in mPFC pyramidal neurons, calculated as the evoked EPSC area divided by the evoked IPSC area evoked by 20 ms laser stimulation as illustrated in (H). KO mice show a selective shift toward excitation in the E/I balance for vHPC, but not callosal, inputs (two-way ANOVA on log-transformed data: genotype effect, F_(1,31)_ = 8.337, p = 0.007, n = 8–10; Sidak’s post hoc test: control vs. KO vHPC, p = 0.025; control vs. KO callosal, p = 0.271). **J,** Input–output slope of fast-spiking interneuron (FSI) responses in vHPC-and callosal-injected control and KO mice. The vHPC-evoked slope is reduced in KO mice, while the callosal-evoked slope is unaffected (vHPC: unpaired t-test on log-transformed data, p = 0.021, n = 8–13; callosal: unpaired t-test, p = 0.731, n = 5–7).

To directly determine whether input-specific recruitment of feedforward inhibition underlies the selective E/I imbalance in pyramidal neurons, we simultaneously recorded optogenetically evoked excitatory synaptic responses from neighboring pyramidal neuron–FSI pairs during stimulation of either vHPC or callosal afferents in control and KO mice (Figure 3A). This paired recording strategy allowed direct comparison of the relative excitatory drive received by neighboring pyramidal neurons and FSIs while controlling for variability in axonal recruitment produced by optical stimulation. In control mice, vHPC and callosal projections preferentially recruited FSIs over neighboring pyramidal neurons, with excitatory responses in FSIs being approximately 2.5–3.3-fold larger than those recorded in PNs (Figure 3B–C, controls). This preferential recruitment was selectively lost for vHPC inputs in KO mice, where excitatory responses in FSIs were markedly reduced, abolishing the normal difference between cell types (Figure 3B, two-way RM ANOVA, genotype × cell type interaction, p = 0.003; Sidak’s post hoc test: control, p < 0.0001; KO, p = 0.822). In contrast, preferential recruitment of FSIs by callosal inputs remained preserved (Figure 3C, cell type effect, p = 0.005). Consistent with this pathway-specific effect, analysis of the PN/FSI EPSP ratio revealed a significant genotype × pathway interaction (two-way ANOVA, p = 0.006), with KO mice differing from controls only for vHPC inputs (Figure 3D, Sidak’s post hoc test, p = 0.005).

**Figure 3.**
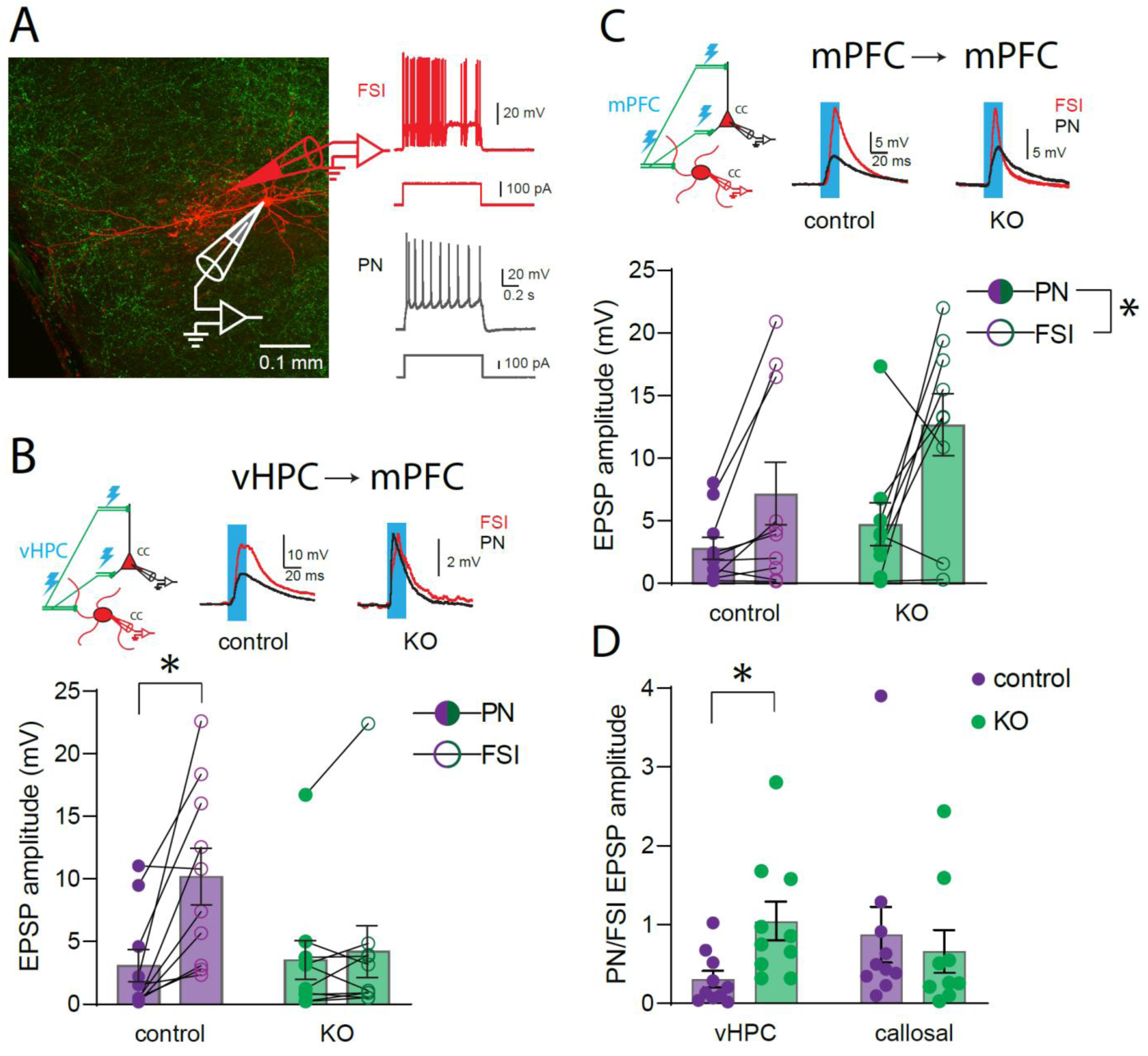
Selective loss of feedforward recruitment of fast-spiking interneurons by ventral hippocampal, but not callosal, inputs in KO mice. **A,** paired whole-cell recording configuration from neighboring a pyramidal neuron (PN) and fast-spiking interneuron (FSI) in mPFC layer 2/3 (left, neurobiotin labeling in red and ChR2 expressing axons in green). Representative firing patterns in response to depolarizing current injection confirm the fast-spiking phenotype of FSIs (red) compared with the regular-spiking pattern of PNs (black). **B,** optogenetic stimulation of ventral hippocampal (vHPC) axons in mPFC (top left, schematic) evokes EPSPs in simultaneously recorded PN–FSI pairs. Representative traces of 447 nm light-evoked EPSPs in FSIs (red) and PNs (black) in control and KO mice. Summary data (bottom) show EPSP amplitude for each PN–FSI pair (connected symbols) in control and KO mice (two-way repeated-measures ANOVA on log transformed data, n = 10: genotype × cell type interaction, F_(1,18)_= 11.98 p = 0.003; Sidak’s post hoc test: control p < 0.0001; KO p = 0.822). **C,** same experimental design as in B, but for optogenetic stimulation of callosal axons (two-way repeated-measures ANOVA on log transformed data, n = 9-10: cell-type effect, F_(1,17)_= 10.33 p = 0.005; genotype effect F_(1,17)_= 1.389 p = 0.255; genotype × cell-type interaction, F_(1,17)_= 0.808 p = 0.381). **D,** summary of the PN/FSI EPSP amplitude ratio for vHPC and callosal inputs in control and KO mice (two-way ANOVA on log transformed data, n = 9-10: genotype × pathway interaction, F_(1,35)_=8.729 p = 0.006; Sidak’s post hoc test for control vs. KO, vHPC p = 0.005, callosal p = 0.575). Data are shown as mean ± SEM, with individual cells/pairs overlaid. *p < 0.05.

In order to study if the synaptic properties are altered in the KO mice, we stimulated the hippocampal and callosal axons with trains of stimulations at 5, 10 and 20 Hz. We found no alteration in the short-term plasticity of vHPC or callosal inputs onto PNs or FSIs at any stimulation frequency tested (Figure 4A-D). The short-term depression was maintained across different laser intensities indicating that synaptic properties of the glutamatergic distal connections seem to be independent of the number of synapses activated. In the pyramidal neurons, two-way ANOVA shows a significant effect between vHPC and callosal synapses (p=0.007), with no effects of genotype (p=0.905) in the magnitude of the short-term depression (Figure 5A). In FSIs on the other hand no differences were observed between genotypes or pathways (Figure 5B). Similar results were obtained when analyzing the pair pulse ratio (Figure 5 C-D). These results indicate that the selective reduction in hippocampal, but not callosal, recruitment of FSIs cannot be explained by alterations in presynaptic glutamatergic release dynamics.

**Figure 4.**
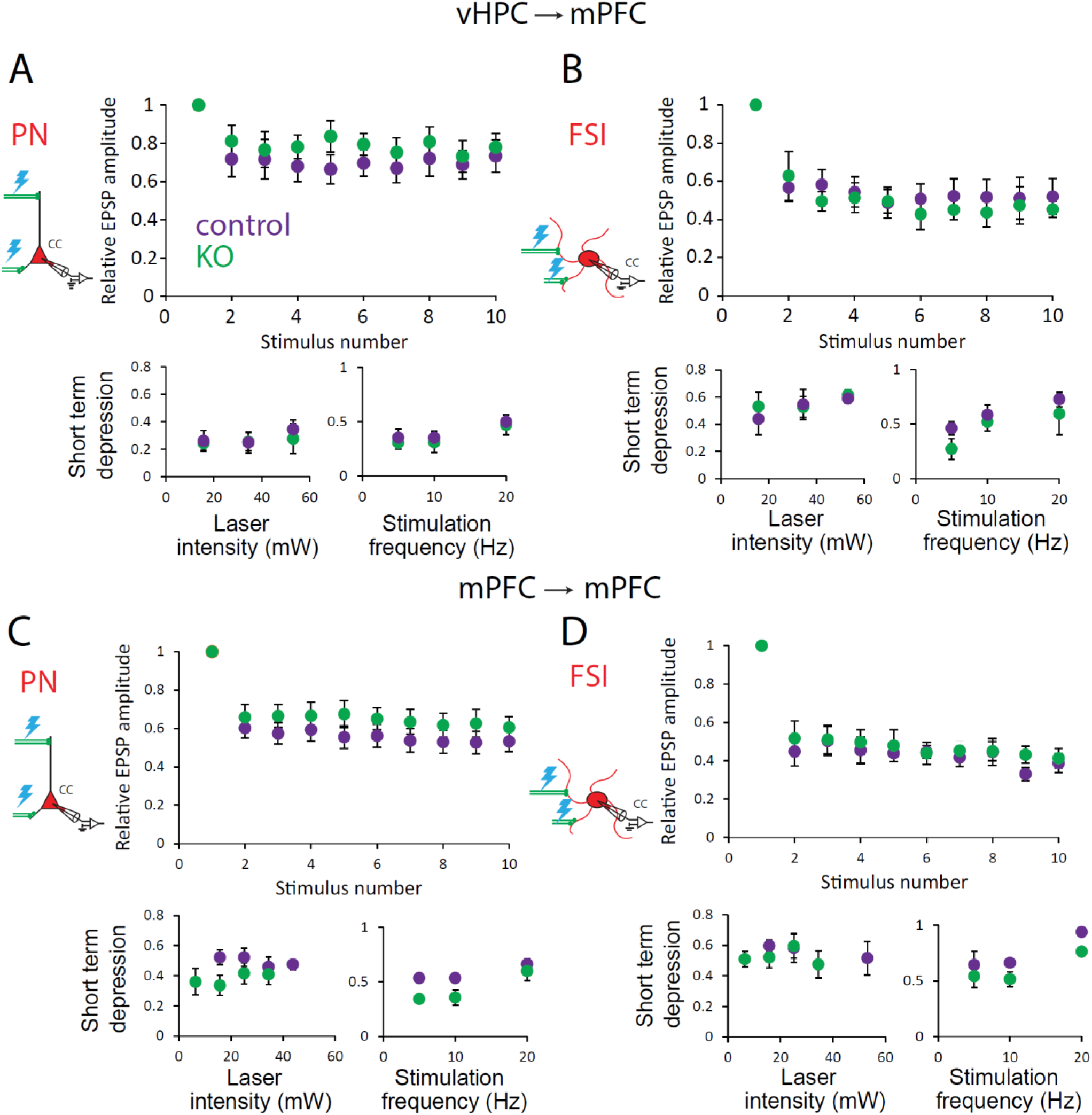
Short-term synaptic plasticity of vHPC and callosal inputs onto mPFC pyramidal neurons and fast-spiking interneurons is unaffected by interneuron NMDAR ablation. Short-term plasticity of vHPC and callosal inputs onto pyramidal neurons (PN) and fast spiking interneurons (FSI). In each case, top, relative EPSP amplitude across consecutive stimuli in a train of 10 stimuli of 20 ms 447nm laser at 10 Hz, in control and KO mice. Bottom, short-term depression as a function of laser stimulation intensity (left), and as a function of stimulation frequency (5, 10, and 20 Hz, right), in control and KO mice. **A,** vHPC inputs onto mPFC PN (mixed-effects ANOVA for laser intensity, n = 9-13: intensity effect F_(2,37)_= 1.617 p = 0.212; genotype effect, F_(1,23)_= 0.042 p = 0.839; intensity x genotype interaction, F_(2,37)_= 0.455 p = 0.638; two way repeated measures ANOVA frequency, n = 9-13: frequency effect F_(2,38)_= 4.822 p = 0.014; genotype effect, F_(1,19)_=0.197 p=0.662; genotype × frequency interaction, F_(2,38)_= 0.012 p = 0.988). **B,** vHPC inputs onto mPFC FSI (two way repeated measures ANOVA for laser intensity, n= 5-9: intensity effect, F_(2,16)_= 0.697 p = 0.513; genotype effect F_(1,8)_= 0.336 p = 0.578; intensity x genotype interaction, F_(2,16)_= 0.105 p = 0.901; two-way repeated measures ANOVA for frequency, n= 5-9: frequency effect F_(2,18)_= 28.88 p < 0.001; genotype effect, F_(1,19)_=0.058 p=0.815; frequency x genotype interaction, F_(2,18)_= 1.024 p = 0.379). **C,** callosal inputs onto mPFC PN (mixed-effects ANOVA for laser intensity, n= 8-11: intensity effect F_(3,26)_= 0.979 p = 0.418; genotype effect, F_(1,19)_= 2.267 p=0.149; intensity x genotype interaction, F_(3,26)_=1.184 p = 0.335; two-way repeated measures ANOVA for frequency, n= 8-11: frequency effect F_(2,30)_= 23.21 p < 0.001; genotype effect, F_(1,15)_= 2.769 p = 0.117; frequency x genotype interaction, F_(2,30)_= 3.032 p = 0.063). **D,** callosal inputs onto mPFC FSI (mixed-effects ANOVA for laser intensity, n = 5-8: intensity effect F_(3,13)_= 1.313 p = 0.312; genotype effect, F_(1,11)_= 0.001 p = 0.973; intensity x genotype interaction, F_(3,13)_= 1.997 p = 0.164; two-way repeated measures ANOVA for frequency, n = 5-8: frequency effect F_(2,20)_= 24.87 p < 0.001; genotype effect, F_(1,10)_= 2.892 p = 0.119; frequency x genotype interaction, F_(2,20)_= 0.288 p = 0.753). Data are shown as mean ± SEM.

**Figure 5.**
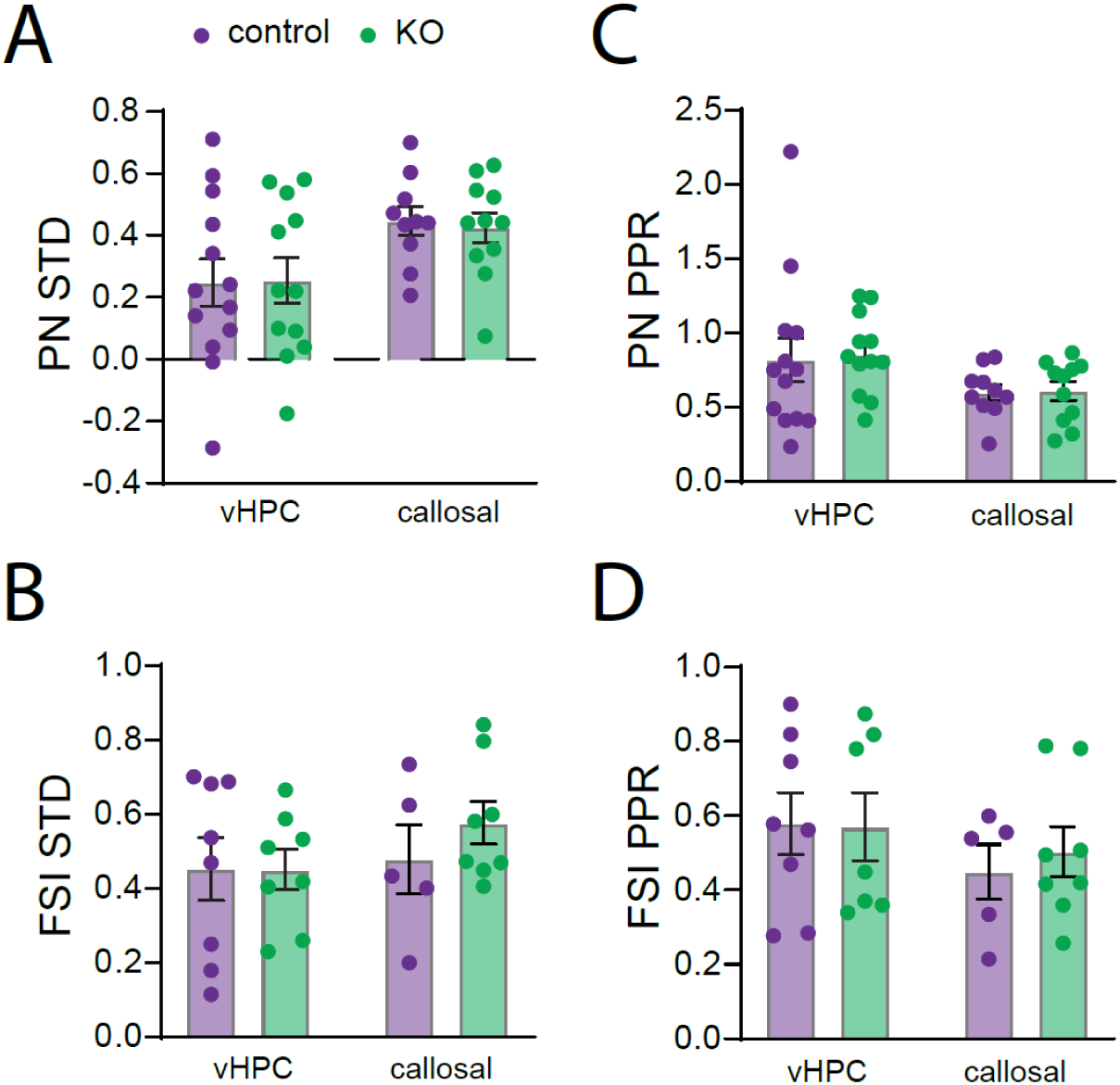
Short-term depression and paired-pulse ratio of vHPC and callosal inputs onto mPFC pyramidal neurons and fast-spiking interneurons are unaffected by interneuron NMDAR ablation. **A,** short-term depression (STD) of vHPC-and callosal-evoked EPSPs onto pyramidal neurons (PN) in control and KO mice (two-way ANOVA, n = 10-13: pathway effect, F_(1,42)_= 7.947 p = 0.007; genotype effect, F_(1,42)_= 0.014 p = 0.905; pathway x genotype interaction, F_(1,42)_= 0.044 p = 0.835). **B,** short-term depression of vHPC-and callosal-evoked EPSPs onto fast-spiking interneurons (FSI) in control and KO mice (two-way ANOVA, n=5-8: pathway effect, F_(1,25)_= 1.079 p = 0.309; genotype effect, F_(1,25)_= 0.438 p = 0.514; pathway x genotype interaction, F_(1,25)_= 0.467 p = 0.501). **C,** paired-pulse ratio (PPR) of vHPC-and callosal-evoked EPSPs onto pyramidal neurons in control and KO mice (two-way ANOVA, n=10-13: pathway effect, F_(1,42)_= 5.315 p= 0.026; genotype effect, F_(1,42)_= 0.051 p = 0.822; pathway x genotype interaction, F_(1,42)_= 0.021 p = 0.885). **D,** paired-pulse ratio of vHPC-and callosal-evoked EPSPs onto fast-spiking interneurons in control and KO mice (two-way ANOVA, n= 5-8: pathway effect, F_(1,24)_= 1.458 p = 0.239; genotype effect, F_(1,24)_= 0.076 p = 0.786; pathway x genotype interaction, F_(1,24)_= 0.151 p = 0.701). Data are shown as mean ± SEM, with individual cells overlaid.

The influence of E/I balance on synaptic integration depends not only on the relative strength of excitation and inhibition but also on their temporal coupling. Feedforward inhibition rapidly truncates EPSPs, thereby limiting the temporal window for synaptic integration. We therefore reasoned that the selective reduction in ventral hippocampal feedforward inhibition in KO mice should prolong hippocampal-evoked EPSPs in pyramidal neurons. To test this hypothesis, we compared the decay kinetics of EPSPs evoked by vHPC and callosal inputs before and after pharmacological blockade of GABA-A receptors. In control mice, blockade of GABA-A receptors with picrotoxin approximately doubled EPSP duration for both vHPC and callosal inputs, demonstrating that feedforward inhibition normally constrains the temporal integration of excitatory inputs (Figure 6A-B, controls). In contrast, in KO mice, picrotoxin failed to significantly prolong EPSPs evoked by vHPC stimulation, indicating that feedforward inhibition recruited by this pathway was already severely compromised. By comparison, picrotoxin continued to markedly prolong callosal-evoked EPSPs in KO mice, consistent with preserved feedforward inhibition in this pathway (Figure 6A-B). Comparison of vHPC responses by two-way repeated-measures ANOVA revealed a significant genotype × treatment interaction (p = 0.016), reflecting the absence of a picrotoxin effect in KO mice (Sidak’s post hoc test: control, p = 0.017; KO, p = 0.994). Consistent with the results shown in figure 3, this deficit was pathway-specific. For callosal inputs, picrotoxin similarly prolonged EPSPs in both control and KO mice (two-way repeated-measures ANOVA, treatment effect, p = 0.0002; Sidak’s post hoc test: control, p = 0.046; KO, p = 0.001), indicating preserved feedforward inhibition. To directly compare the contribution of feedforward inhibition across pathways, we quantified the picrotoxin/ACSF EPSP decay ratio (Figure 6C). This analysis revealed a significant genotype × pathway interaction (two-way ANOVA, p = 0.026), with a selective reduction in the effect of picrotoxin on vHPC inputs in KO mice (Sidak’s post hoc test, p = 0.023). Together, these findings demonstrate that the input-specific reduction in recruitment of feedforward inhibitory circuits selectively prolongs hippocampal synaptic integration in mPFC pyramidal neurons.

**Figure 6.**
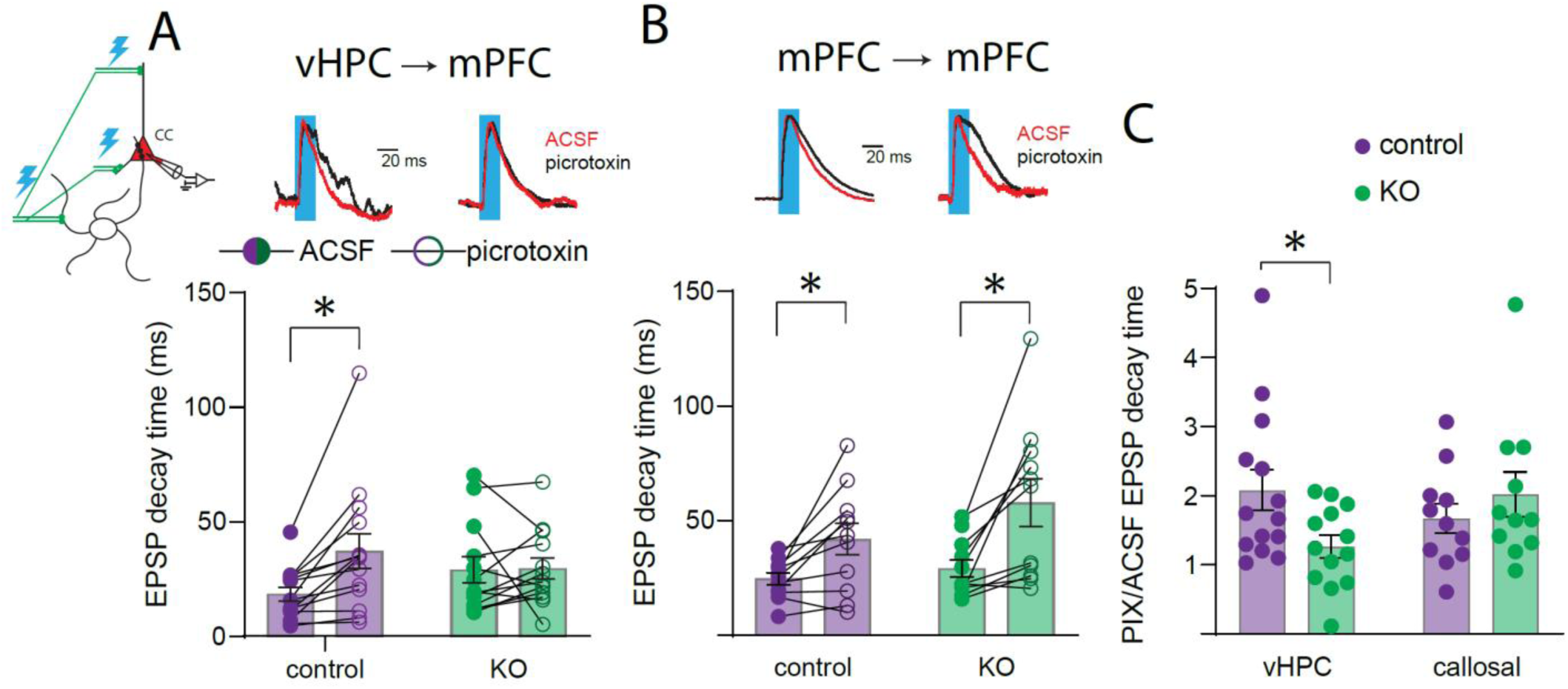
Selective reduction of feedforward inhibition of vHPC inputs onto apical dendrites of mPFC pyramidal neurons following interneuron NMDAR ablation. **A,** schematic of recording configuration (top left) and representative EPSPs evoked by optogenetic stimulation of ventral hippocampal (vHPC) axons before (ACSF, red) and after blockade of GABA_A receptors with picrotoxin (black). **Bottom,** EPSP decay time for individual pyramidal neurons recorded in ACSF and picrotoxin in control and KO mice (two-way repeated-measures ANOVA, n=13-14: genotype × treatment interaction, F_(1,25)_=6.594 p = 0.016; Sidak’s post hoc test: control, p = 0.017; KO, p = 0.994). **B,** same as A, for optogenetic stimulation of callosal axons (two-way repeated-measures ANOVA, n=11: treatment effect, F_(1,20)_= 21.24 p < 0.001, genotype effect, F_(1,20)_= 1.658 p = 0.213; genotype × treatment interaction, F_(1,20)_= 1.306 p = 0.267; Sidak’s post hoc test: control, p = 0.046; KO, p = 0.001). **C,** ratio of EPSP decay time in picrotoxin relative to ACSF (PIX/ACSF) for vHPC and callosal inputs in control and KO mice (two-way ANOVA, n=11-14: genotype × pathway interaction, F_(1,45)_= 5.288 p = 0.026; Sidak’s post hoc test for control vs. KO: vHPC, p = 0.023; callosal, p = 0.723). Data are shown as mean ± SEM, with individual cells overlaid (paired data connected by lines in A and B). *p < 0.05.

## Discussion

The present study identifies pathway-specific disruption of excitatory/inhibitory balance as a previously unrecognized consequence of interneuron NMDAR hypofunction. By combining structural analysis with pathway-specific optogenetics and electrophysiology, we demonstrate that both structural and functional E/I imbalance emerge selectively at ventral hippocampal, but not callosal, inputs onto medial prefrontal pyramidal neurons. This selective imbalance results from impaired recruitment of feedforward inhibition, leading to prolonged temporal integration of hippocampal inputs. These findings extend the current view of cortical E/I imbalance from the net balance of excitation and inhibition received by a neuron to a pathway-specific property of individual inputs, proposing impaired inhibitory filtering as the mechanism by which interneuron dysfunction selectively biases cortical information processing. More broadly, our results indicate that measurements of the net neuronal E/I balance may obscure substantial pathway-specific alterations, highlighting the importance of resolving excitation and inhibition at the level of defined synaptic circuits.

Our findings also raise the possibility that pathway-specific E/I imbalance may be further constrained by dendritic compartment. Pyramidal neurons integrate bottom-up and feedforward inputs, such as thalamic and lower-order cortical projections, primarily within basal dendrites, whereas long-range feedback projections from higher cortical areas preferentially target the apical tuft^39–41^. Because apical and basal dendritic compartments are electrotonically and functionally segregated, they can independently integrate distinct streams of synaptic information^42^. In this context, selective disruption of inhibitory control within a particular dendritic domain would be expected to differentially affect the computations supported by the afferents terminating in that compartment. Our finding that structural and functional E/I imbalance selectively affects ventral hippocampal inputs onto the apical dendritic arbor suggests that interneuron dysfunction may distort cortical processing not only in an input-specific manner but also according to the compartmental organization of pyramidal neurons. Such compartment-specific disruption could further increase the computational diversity of circuit dysfunction by selectively altering the integration of distinct classes of cortical information. Thus, rather than representing a uniform property of cortical circuits, E/I imbalance may be organized along two complementary dimensions: the identity of the afferent pathway and the dendritic compartment in which that pathway is integrated. This organization may allow distinct afferent pathways to independently establish and maintain their own E/I set points.

If E/I balance is independently regulated across afferent pathways and dendritic compartments, an important question is how such selective organization influences cortical computations. Previous studies have proposed that cortical E/I imbalance contributes to the reduced signal-to-noise ratio observed in schizophrenia, leading to impaired neuronal tuning and inefficient information processing^43,44^. Our findings extend this framework by suggesting that reduced signal-to-noise may not be a global property of cortical networks but instead emerge selectively within individual afferent pathways. Feedforward inhibition is thought to improve the signal-to-noise ratio by rapidly suppressing excitatory responses, narrowing the temporal window for synaptic integration, and limiting the influence of weak or asynchronous inputs on spike generation. Our findings suggest that this inhibitory filtering can be independently disrupted across convergent afferent pathways. For example, the selective loss of feedforward inhibition at ventral hippocampal synapses prolongs hippocampal-evoked depolarization, biasing temporal integration in favor of hippocampal activity over competing afferents. From this perspective, the proposed reduction in signal-to-noise ratio for schizophrenia and other disorders, characterized by E/I imbalance, may arise not only from a generalized increase in cortical noise, but also from selective failures of inhibitory filtering affecting subsets of synaptic inputs. Such pathway-specific defects would prolong the temporal influence of selected afferents, biasing their contribution to spike generation while diminishing the relative impact of competing inputs. At the level of neuronal ensembles, this imbalance would distort cortical representations by preferentially recruiting activity patterns driven by the affected pathways while underrepresenting information conveyed by preserved inputs. Consequently, cortical processing would become progressively biased toward selected sources of information, reducing the fidelity of cortical representations without requiring a generalized increase in cortical excitability.

Activity-dependent plasticity is a fundamental property of excitatory synapses and is typically expressed in an input-specific manner. Hebbian and homeostatic mechanisms allow individual afferent pathways to be independently strengthened or weakened during development and experience. Our findings raise the possibility that inhibitory circuits are refined according to the same principle, allowing feedforward inhibition to establish pathway-specific E/I balance for individual afferent systems.

Although demonstrated here in a single model of interneuron NMDAR hypofunction, we propose that input-specific E/I imbalance may represent a broader organizing principle of circuit dysfunction following interneuron impairment. In this model, selectively impaired feedforward inhibitory filtering represents one mechanism by which this selective imbalance emerges. Given the remarkable diversity of cortical interneuron subtypes and their specialized roles in regulating distinct dendritic compartments, afferent pathways, and circuit computations^45–48^, dysfunction of different interneuron populations may generate distinct signatures of input-specific E/I imbalance. Thus, this framework may extend beyond schizophrenia to other disorders associated with E/I imbalance, including autism spectrum disorders, where selective dysfunction of defined circuits is increasingly recognized. Future studies should establish the extent to which input-specific E/I imbalance represents a general principle of circuit pathology across brain regions and disease models.

## Acknowledgments

We thank Analia Lopez Diaz, Agostina Presta, Veronica Risso, and Lucia Garbini for technical assistance, and the GNS group for feedback on previous drafts.

## Author Contributions

CAPA, JEB & DEP designed the study. CAPA & DEP performed the experiments and collected data. CAPA & DEP analyzed the data. JEB & DEP wrote the manuscript. CAPA, JEB & DEP reviewed and edited the manuscript.

## Funding

This work was supported in part by Agencia Nacional de Promoción Científica y Tecnológica, Universidad de Buenos Aires, and CONICET. All authors were supported by the Consejo Nacional de Investigaciones Científicas y Técnicas (CONICET) and the Universidad de Buenos Aires, Argentina.

## Competing Interests

The authors declare no competing interests.

## Notes

### Competing Interest Statement

The authors have declared no competing interest.

### Summary of Updates

we updated figure 2, corrected the pseudocolor LUT of bright field in image B, and removed bright field for image D. We rearranged the panels of figure 3, and figure 6 We summarized the materials and methods section in the ms and included a full description in the supplemental materials and methods section

