## supplement for "NMDA receptor hypofunction on GABAergic interneurons results in input-specific Excitatory/Inhibitory imbalance in pyramidal neurons of medial prefrontal cortex"

**Running title:** NMDAr interneuron hypofunction leads to input-specific E/I imbalance in mPFC

**Corresponding authors:**

Juan E. Belforte, 2155 Paraguay[st] 7 floor, Ciudad de Buenos Aires, Argentina (1121), +54-11-5285-3309,.

Diego E Pafundo, 2155 Paraguay St, 7 floor, Buenos Aires (1121), Argentina. +54-11-5285-3312.

### Supplemental Materials and Methods

#### Animals

Male and female Ppp1r2-Cre/homozygously-floxed-Grin1 knockout mice (Grin1 KO, mutant or KO mice) and Cre-negative control littermates (homozygously-floxed-Grin1: control) group-housed on a 12:12 light/dark cycle with ad libitum access to food and water were used following protocols approved by the University of Buenos Aires School of Medicine IACUC and government regulations (SENASA, Argentina). The KO mice has already been extensively characterized regarding the Cre-mediated expression and recombination pattern in different brain regions, including mPFC and Hippocampus[31,32]. In these mice, genetic deletion of obligatory Grin1 subunit occurs in ~50% of cortical and hippocampal GABAergic interneurons, including over 75 percent of parvalbumin positive neurons, from the second postnatal week (KO mice). Stable Cre-mediated recombination was previously observed from postnatal week 4 to 24. All mice were maintained in C57BL/6 background.

#### Surgery and viral labeling

Mice (8-10 weeks) were anaesthetized using isoflurane (2.5% for induction and 0.5-1.0% for maintenance, in 95% oxygen) and mounted in a stereotaxic frame (David Kopf Instruments) with a heating pad to maintain body temperature. An incision was made to expose the skull for stereotaxic alignment using bregma and lambda and a borehole was either drilled to target the vHPC (-3.3 mm AP, -+/-3.3mm ML relative to bregma) or the mPFC (+1.9 mm AP, +/-0.3- +/-0.5mm ML). An adeno-associated virus (AAV) vector tagged with enhanced yellow fluorescent protein (eYFP) and expressing channelrhodopsin (ChR2) under the CaMKII promoter (AAV-CaMKIIa-hChR2(H134R)-EYFP, 5.1 1012 vg ml-1, cat # 26969 Addgene, USA) was administered by intracerebral injection using pipettes pulled from borosilicate glass at 0.1 µl min-1. For targeting the vHPC 0.5 µl of virus was microinjected at -3.2mm and -2.7mm DV from the surface of the brain and for the mPFC 0.5 µl was microinjected at -1.9 mm DV from the surface of the brain. After injection, the pipette was kept at the injection site for 5-10 min before being slowly withdrawn. The scalp was then sutured and 5mg kg-1 flunixin was subcutaneously administered.

The areas of the ventral hippocampus and the contralateral mPFC transduced by the AAV virus were estimated with immunohistochemistry by measuring the area of EYFP fluorescence in the CA1 region of the coronal plane with the largest transduction of EYFP and the septal-temporal span of infection was estimated by counting the number of continuous coronal planes containing CA1 labeled with EYFP and the area of the contralateral mPFC transduced was estimated by measuring the area of EYFP fluorescence in the mPFC cortical layers and rostro caudal infection was estimated by counting the number of continuous coronal planes containing labeled mPFC with EYFP.

For immunohistochemistry experiments, immediately after the stereotaxic surgery 50 µl of an AAV-PHP.eB adeno-associated virus, AAV PHP.eB -pAAV-CAG-tdTomato (cat # 59462-PHPeB, 1.1 1011 vg ml-1) was administered by retro-orbital injection to sparsely label neurons in the mPFC using a 30-gauge syringe under isoflurane anesthesia.

### Immunohistochemistry

The AAV-PHP.eB adeno-associated virus sparsely labeled neurons with tdTomato and the AAV-CaMKIIa-ChR2-EYFP virus traced the pyramidal neuron long-range projections from either the vHPC and the contralateral mPFC. 30-50 days post-surgery, animals were transcardially perfused and 30 µm coronal slices were obtained with a freezing microtome (SM2010 Leica Microsystems) and processed for immunofluorescent staining. Slices containing sparse labelled pyramidal neurons were rinsed at RT (3 times for 10min each) in PBS and blocked at RT with 0.3% Triton X-100, 5% normal goat serum (NGS) and 1% BSA for 2 hs. Slices were then incubated overnight at 4 °C with 0.1% Triton X-100, 1% NGS and monoclonal anti-mCherry (rabbit 1:1000, ab167453, abcam), anti-GFP (chicken 1:2000, GFP-1010 AvesLabs) and anti-VGAT (guinea pig 1:600, 131 004, Synaptic Systems) primary antibodies. Slices were then rinsed trice in PBS (10 min each) at room temperature and incubated with PBS containing 0.1% Triton X-100, 1% NGS, Cy3 goat anti-rabbit (1:500; 111-165-144, Jackson ImmunoResearch Inc.), biotinylated goat anti-guinea pig (1:500, BA-7000, Vector Laboratories) and AlexaFluor 488 goat anti-chicken (1:1000, A11039 Thermo Fisher) antibodies for 2 h at room temperature. After rinsing 3 times for 10’ in PBS, slices were incubated in PBS containing streptavidin-Alexa Fluor 647 (1:500, 016-600-084 Jackson ImmunoResearch Inc.). Slices were then rinsed twice in PBS (10 min each) at room temperature and mounted with vectashield (H1000, Vector Laboratories). Images were acquired with a Zeiss Axio Imager.M2 with Apotome.2-Neurolucida Stereoinvestigator system (MBF, USA). Pyramidal neuron identification and determination of dendrite order was performed under a 20x objective (20 planes, 1.5 µm z-interval) whereas dendrite sections, spines, YFP boutons and VGAT puncta were acquired under 63x objective (75-100 planes, 0.2 µm z-interval). Only td-Tomato labeled pyramidal neurons where the basal dendrites could be traced to the soma, and the apical tuft dendrites could be traced to the apical dendrite in a mPFC region with YFP axon afferents and sparse neuron labeling were used. Dendritic segments of 2nd to 4th order, spines and YFP and VGAT puncta were identified and counted from the images acquired using Neurolucida (MBF, USA), YFP and VGAT puncta were counted as appositions when they were in direct contact with the dendritic spines and dendrite shaft respectively.

### Slice physiology

Experiments were conducted in brain slices prepared from the mPFC of 21-24 weeks-old mice. These mice were injected with AAV particles 13-14 weeks prior to experiments. Mice were deeply anaesthetized with isoflurane and decapitated. Brains were quickly removed and immersed in ice-cold slicing solution containing (in mM): 210 sucrose, 10 NaCl, 1.9 KCl, 1.2 $Na_2HPO_4$, 33 $NaHCO_3$, 6 $MgCl_2$, 1 $CaCl_2$ and 10 glucose; pH 7.3-7.4 when bubbled with 95% $O_2$ and 5% $CO_2$. 300 µm coronal slices of the mPFC and the ventral hippocampus were sectioned using a vibrating microtome (Pelco 1000, Ted Pella, Inc, USA). Slices containing the ventral hippocampus were fixed in paraformaldehyde (4% in PBS) and mounted for YFP fluorescence images, and slices containing the mPFC were immediately placed in an incubation chamber filled with artificial cerebrospinal fluid (ACSF) maintained at 36°C and containing (in mM): 125 NaCl, 2.5 KCl, 1.25 $Na_2HPO_4$, 10 glucose, 25 $NaHCO_3$, 0.4 ascorbate, 1 $MgCl_2$ and 2 $CaCl_2$, pH 7.3-7.4 when gassed with 95% $O_2$ and 5% $CO_2$. After 5 min incubation at 36°C, brain slices were stabilized at room temperature in the same solution for at least 30 min before they were transferred to the recording chamber. For recordings, slices were transferred to a submersion chamber and superfused at 2 ml min−1 with oxygenated ACSF at 30-32°C. Whole-cell recordings were obtained from visually identified pyramidal neurons and fast spiking interneurons consistent with parvalbumin positive neurons in layer 2-3 of the mPFC using a Nikon microscope equipped with IR-DIC optics as described before[30,33,34]. Pipettes pulled from borosilicate glass had a resistance of 4-6 MΩ when filled with the intracellular solution (in mM): 120 potassium gluconate, 10 KCl, 10 HEPES, 0.2 EGTA, 4.5 MgATP, 0.3 NaGTP, 14 sodium phosphocreatine, 0.1-0.2% neurobiotin; pH adjusted to 7.2-7.4 with KOH. Recordings were obtained using Multiclamp 700B amplifiers (Molecular Devices, USA). Signals were low-pass filtered at 6 kHz and digitized at 20 kHz using DigiData 1200 acquisition interfaces (Molecular Devices, USA). Data acquisition and analysis were performed using Clampex 10.2 and ClampFit 10.2 software (Molecular Devices, USA). Only recordings with a stable series resistance of <20 MΩ were used for analysis. In the current clamp, series resistance and pipette capacitance were canceled using bridge and capacitance neutralization. Neurons included in this study had resting membrane potential between -60 and -80 mV. EPSCs, IPSCs and EPSPs were elicited in pyramidal neurons, and EPSCs and EPSPs in fast spiking interneurons, by stimulating the ChR2 expressing vHPC and contralateral mPFC terminals with 447nm laser (Tolket, Argentina) through a water immersion 40X microscope objective (6-53 mW in the focal plane of the objective) centered in the recorded neuron. In the experiments shown in Fig 6, evoked EPSP responses were analyzed in the absence and presence of the GABAA receptor blocker picrotoxin (P1675, Merck). After recording the evoked EPSCs in ACSF, slices were incubated with 50 µM picrotoxin for 10 min and then evoked EPSCs in the presence of picrotoxin were recorded.

After recordings, the slices were quickly fixed with paraformaldehyde (4% in PBS), and recorded neurons were imaged using fluorescence labeled streptavidin. The recorded slices incubated with streptavidin-Cy3 (1:500 Invitrogen), 0.5% Triton X-100 in 0.1M PBS overnight at 4°C. The sections were rinsed with PBS and mounted with Vectashield (H-1000 Vector Laboratories). Image stacks of labeled neurons were acquired with an epifluorescence microscope (Axio Imager.M2, Zeiss) equipped with structured light illumination (Apotome.2, Zeiss).

Quantification and Statistical Analysis

Structural E/I was calculated as the density of excitatory inputs (total as the density of spines or specific as the density of vHPC or callosal appositions to dendritic spines) divided by the density of VGAT appositions to the dendritic shaft. Functional E/I was calculated as the area of the evoked EPSC divided by the area of the evoked IPSC, recorded at -60mV and 0mV respectively. In experiments of Figs 4 and 5, short term depression was calculated from the trains of ChR stimulation as 1- B/A, where B is the average of the amplitude of the last 4 evoked EPSPs in the stimulation train and A is the amplitude of the first evoked EPSP in the train[33]. From the same experiments, pair pulse ratio was calculated from the amplitudes of the second and first evoked EPSPs in the train.

Results are expressed as mean ± SEM, and all bar plots are shown along individual data. Normality of data distributions was estimated with Shapiro-Wilk tests in all experiments. Significance between group means was determined using paired or two sample t-tests, or ANOVA, as indicated in each case. When normality was not met, data was log transformed and significance between groups was determined using the log-transformed data as indicated in each case. Significant p values are indicated along with statistical parameters such as number of cells and animals used in each experiment in the Results section and Figure Legends. No animals that fulfill our quality inclusion criteria were excluded from the analysis and only one neuron was excluded from the structural E/I balance analysis (figure 1I and Suppl figure 3D) because it was identified as an outlier by the Grubbs test (alpha=0.05). Analyses were conducted in Prism (Graphpad) and Statistica (StatSoft).

Supplemental table

| | vHPV→mPFC | | | mPFC→mPFC | | |
|---|---|---|---|---|---|---|
| | Control (n=9) | KO (n=10) | p | Control (n=8) | KO (n=8) | p |
| EPSC delay (ms) | 2.688±0.171 | 4.224±0.790 | 0.089 | 3.482±0.384 | 3.302±0.739 | 0.832 |
| IPSC delay (ms) | 6.605±1.579 | 8.999±1.874 | 0.348 | 7.791±0.749 | 8.646±1.962 | 0.689 |
| Δ delay (IPSC -EPSC ms) | 4.139±1.680 | 4.776±1.831 | 0.731 | 4.308±0.627 | 5.345±2.064 | 0.638 |
| EPSC peak delay (ms) | 7.056±1.247 | 10.424±1.480 | 0.104 | 9.867±1.517 | 7.535±1.331 | 0.267 |
| EPSC area (pA $s^{-1}$) | 0.569±0.229 | 1.603±0.979 | 0.342 | 0.494±0.192 | 2.092±0.906 | 0.106 |
| IPSC area (pA $s^{-1}$) | 1.218±0.219 | 1.101±0.570 | 0.856 | 0.506±0.112 | 1.928±0.809 | 0.225 |

Supplemental Table 1. properties of vHPC and callosal evoked EPSC and IPSC on pyramidal neurons. Data are mean ± SEM, p values from unpaired t test between control and KO.

Supplemental Figure 1

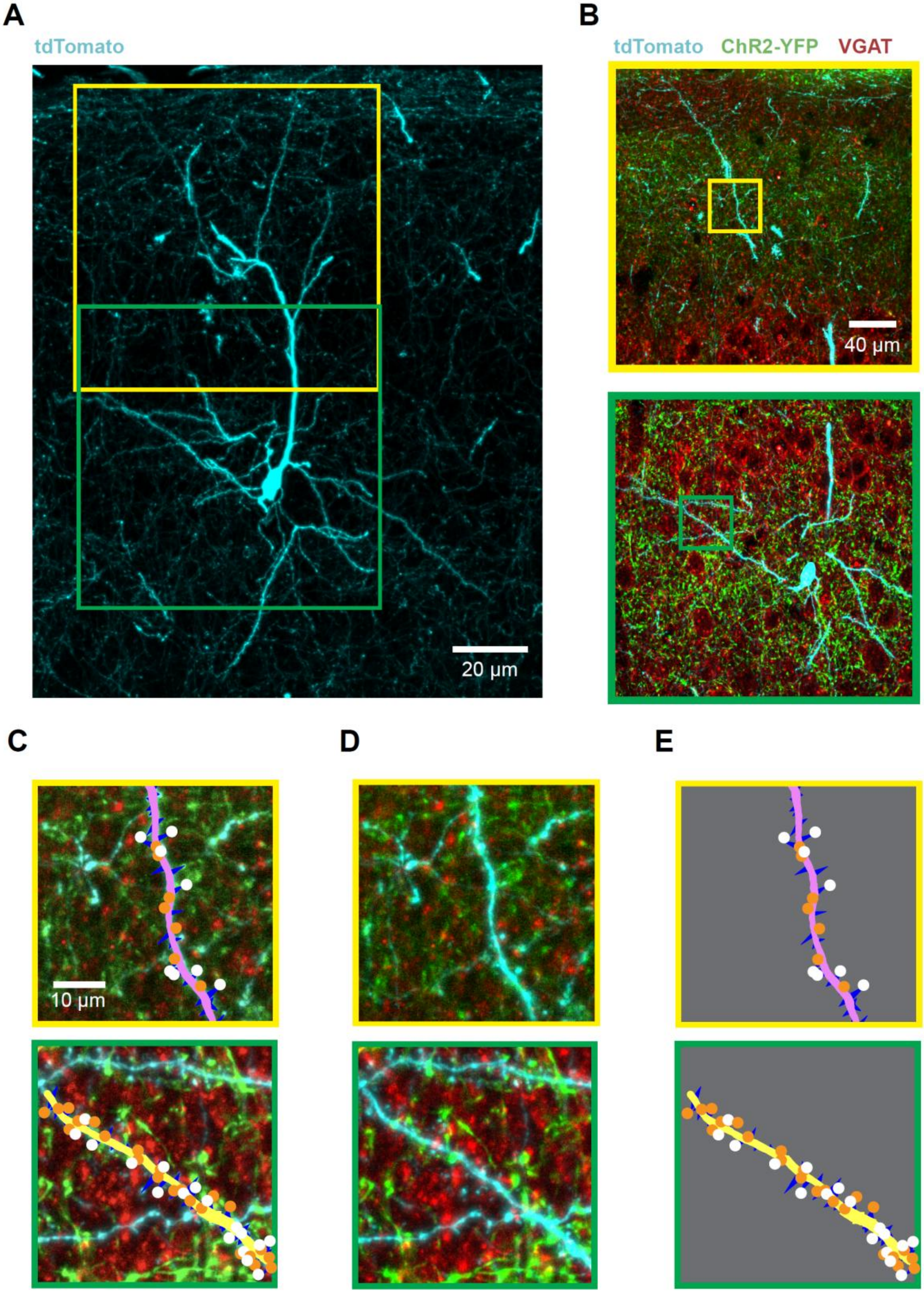

**Supplementary Figure 1. Reconstruction of apical and basal dendrites and identification of excitatory and inhibitory contacts for the analysis of the structural E/I ratio from mPFC pyramidal neurons. A.** Z-projection of a fluorescence image stack from a tdTomato labeled (cyan) pyramidal neuron from the mPFC. The yellow square indicates the location of the fluorescence image stack obtained from an apical dendrite, while the green square indicates that of a basal dendrite. **B.** Z-projection of fluorescence image stacks obtained from the dendrites indicated in A. **Top,** apical dendrite. The yellow square indicates the segment of apical dendrite detailed in C-E top panels. **Bottom,** basal dendrite. The green square indicates the segment of basal dendrite detailed in C-E bottom panels. **C.** Details of the fluorescence images from B, showing examples of reconstructed dendrites. The dendritic shafts with their corresponding spines are superimposed to the fluorescence images. White circles indicate detected YFP+ excitatory inputs, while orange circles indicate VGAT+ inhibitory inputs. **Top,** apical dendrite. Dendritic shaft shown in pink. **Bottom,** basal dendrite. Dendritic shaft shown in yellow. **D.** Z-projection of fluorescence image stacks from C. **Top,** apical dendrite. **Bottom,** basal dendrite. **E.** Images from the reconstructed dendritic shafts with their spines, and excitatory and inhibitory inputs. **Top,** apical dendrite. **Bottom,** basal dendrite.

Supplemental Figure 2

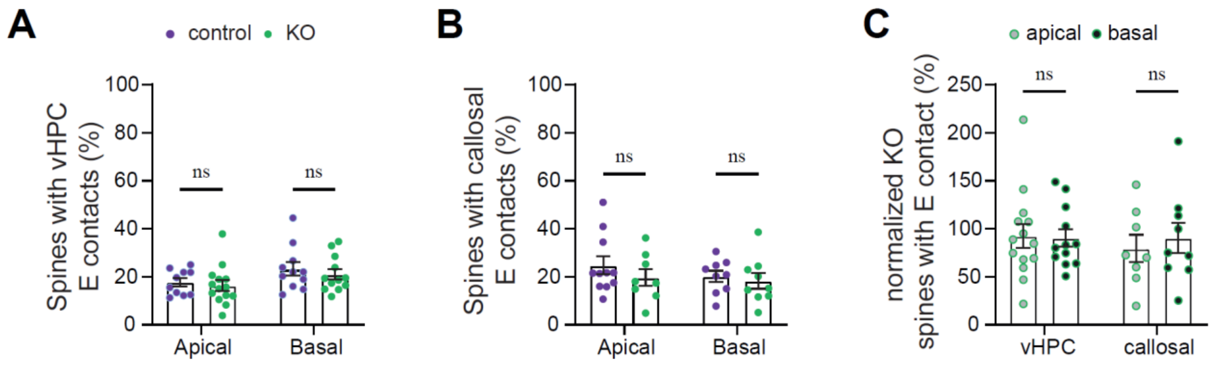


**Supplementary Figure 2. Dendritic spines from mPFC pyramidal neurons receiving either long-range vHPC or contralateral callosal mPFC excitatory projections. A,** dendritic spines receiving excitatory (E) contacts from long-range vHPC projections. Percentage of total dendritic spines from apical and basal dendrites of mPFC pyramidal neurons in control and KO mice (mixed-effects ANOVA, n=10-14 dendrite compartment factor, $F_{(1,20)}$=10.43 p = 0.0042). **B,** dendritic spines receiving excitatory (E) contacts from contralateral callosal mPFC projections. Percentage of total dendritic spines from apical and basal dendrites of mPFC pyramidal neurons in control and KO mice (mixed-effects ANOVA, n=8-11 dendrite compartment factor, $F_{(1,20)}$=10.43 p = 0.0042; genotype factor, $F_{(1,23)}$=0. 2856 p = 0.5982; interaction genotype × dendrite compartment, $F_{(1,20)}$=0.0484 p = 0.8281, n.s.). **C,** dendritic spines from mPFC pyramidal neurons receiving either long-range vHPC or contralateral callosal mPFC excitatory projections, normalized to the respective control group (two way ANOVA, n=8-14 dendrite compartment factor $F_{(1,15)}$=0. 2216 p = 0.6446; genotype factor, $F_{(1,24)}$=0. 4823 p = 0.8280; interaction genotype × dendrite compartment, $F_{(1,15)}$=0.1675 p = 0.6881, n.s.). In all cases, data are shown as mean ± SEM with individual dendritic segments overlaid.

Supplemental Figure 3

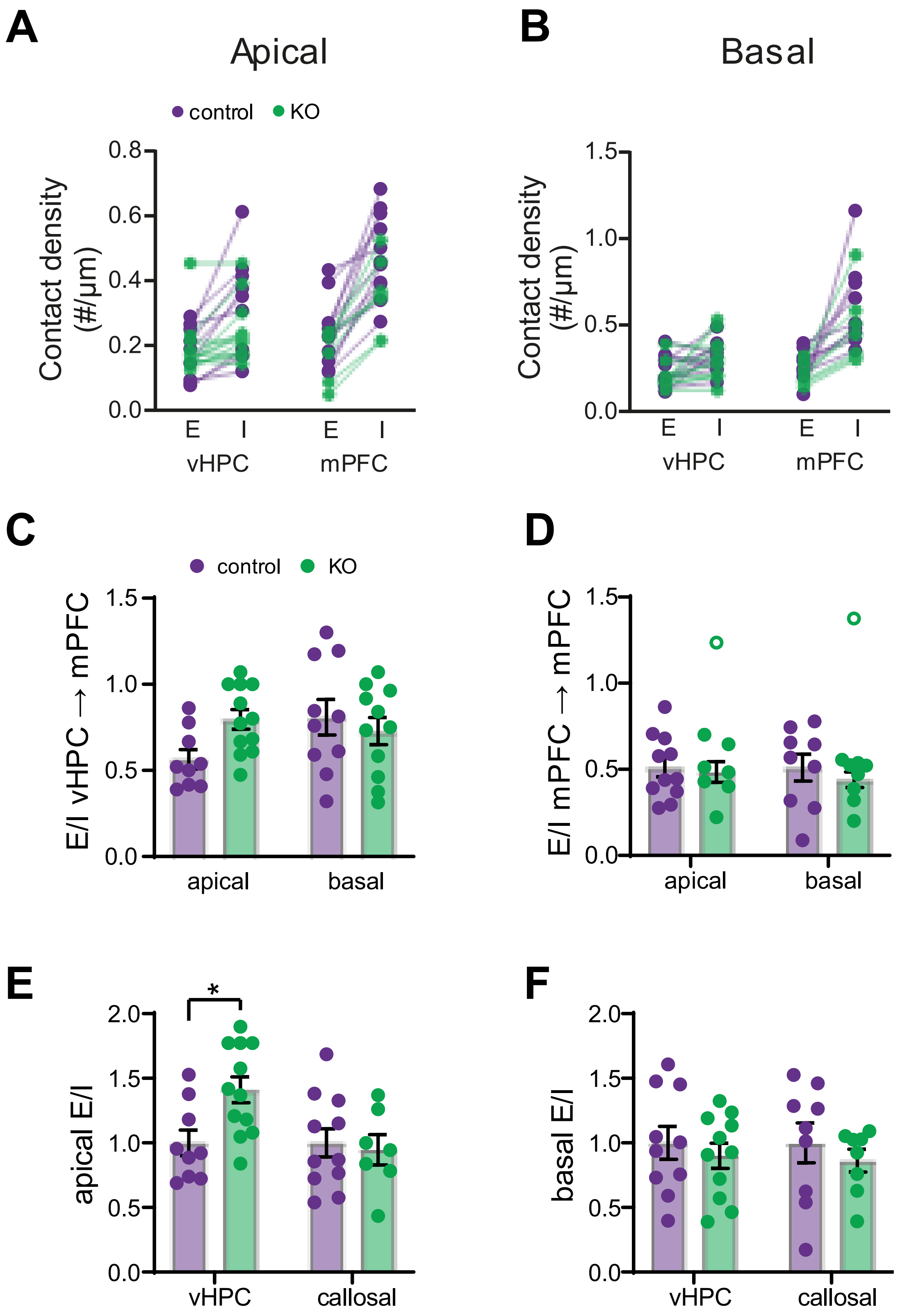

**Supplementary Figure 3. Structural E/I ratio of individual afferent pathways onto mPFC pyramidal neuron dendrites, related to Figure 1. A-B,** density of vHPC and callosal appositions to dendritic spines (E) and paired density of VGAT appositions to the same dendrites (I) for apical (**A**) and basal dendrites (**B**). **C,** non-normalized structural E/I ratio of ventral hippocampal (vHPC) inputs onto apical and basal dendrites of mPFC pyramidal neurons in control and KO mice (mixed-effects ANOVA, n=9-12 genotype × dendrite compartment, $F_{(1,16)}=4.592$ $p = 0.048$). **D,** non-normalized structural E/I ratio of contralateral mPFC (callosal) inputs onto apical and basal dendrites of mPFC pyramidal neurons in control and KO mice (mixed-effects ANOVA, n=7-11 dendrite compartment effect, $F_{(1,13)}=0.211$ $p = 0.653$; genotype effect, $F_{(1,18)}=0.278$ $p = 0.604$; dendrite compartment × genotype interaction, $F_{(1,13)}=0.003$ $p = 0.955$; one data was not included in the ANOVA in the KO apical and basal since it was identified as an outlier, Grubbs method alpha=0.05, and is shown as an open circle). **E,** structural E/I ratio in the apical dendritic compartment for vHPC and callosal inputs, normalized to the respective control group (two way ANOVA, n=7-12 input region × genotype interaction, $F_{(1,35)}=4.465$ $p=0.042$, Sidak's post hoc test: vHPC, $p = 0.0174$; callosal, $p = 0.935$). **F,** structural E/I ratio in the basal dendritic compartment for vHPC and callosal inputs, normalized to the respective control group (two way ANOVA, n=7-11 input region $F_{(1,34)}=0.025$ $p=0.876$, genotype, $F_{(1,34)}=0.966$ $p=0.333$, input region × genotype interaction, $F_{(1,34)}=0.025$ $p=0.876$). Data are shown as mean ± SEM with individual dendritic segments/neurons overlaid. *$p < 0.05$.
